# Reference Set of *Mycobacterium tuberculosis* Clinical Strains: A tool for research and product development

**DOI:** 10.1101/399709

**Authors:** Sonia Borrell, Andrej Trauner, Daniela Brites, Leen Rigouts, Chloe Loiseau, Mireia Coscolla, Stefan Niemann, Bouke De Jong, Dorothy Yeboah-Manu, Midori Kato-Maeda, Julia Feldmann, Miriam Reinhard, Christian Beisel, Sebastien Gagneux

## Abstract

The *Mycobacterium tuberculosis* complex (MTBC) causes tuberculosis (TB) in humans and various other mammals. The human-adapted members of the MTBC comprise seven phylogenetic lineages that differ in their geographical distribution. There is growing evidence that this phylogenetic diversity modulates the outcome of TB infection and disease. For decades, TB research and development has focused on the two canonical MTBC reference strains H37Rv and Erdman, both of which belong to Lineage 4. Relying on only a few laboratory-adapted strains can be misleading as study results might not be directly transferrable to clinical settings where patients are infected with a diverse array of strains, including drug-resistant variants. Here, we argue for the need to expand TB research and development by incorporating the phylogenetic diversity of the MTBC. To facilitate such work, we have assembled a group of 20 genetically well-characterized clinical strains representing the seven known human-adapted MTBC lineages. With the “MTBC clinical strains reference set” we aim to provide a standardized resource for the TB community. We hope it will enable more direct comparisons between studies that explore the physiology of MTBC beyond the lab strains used thus far. We anticipate that detailed phenotypic analyses of this reference strain set will increase our understanding of TB biology and assist in the development of new control tools that are universally effective.

## Introduction

Tuberculosis (TB) remains an urgent public health problem causing 10.4 million new cases and 1.7 million deaths every year. The TB epidemic is worsening due to growing drug resistance and the absence of a universally effective vaccine against the transmissible pulmonary form of the disease (1).

The outcome of TB infection and disease is highly variable, ranging from rapid clearing by the innate immune response to life-long latent infection and various forms of active pulmonary and extrapulmonary disease. In the past, most of this variation has been attributed to the host and environmental factors. Because of the limited genetic diversity within the *Mycobacterium tuberculosis* complex (MTBC) compared to other bacteria (2), the view has been that no relevant phenotypic variation should be expected. However, recent advances in whole genome sequencing of large MTBC clinical strain collections from global sources have revealed more genomic diversity than previously appreciated. Specifically, the human-adapted MTBC comprises seven phylogenetic lineages that differ in their geographic distribution, and individual members of these lineages can differ by up to ∼2,000 single nucleotide polymorphisms (SNPs). This is equivalent to the phylogenetic distance between *M. tuberculosis* sensu stricto and *M. bovis*, which is a typical pathogen of cattle.

In addition to the genomic diversity across MTBC clinical strains, findings from many experimental studies have led to a change in paradigm by demonstrating the phenotypic impact of this genotypic diversity. For example, studies have reported differences between clinical strains in their transcriptomic profile (3, 4), protein and metabolite levels (5) (4), methylation profiles (5), drug susceptibility (6) and cell wall structure (7-9). In addition, MTBC genetic diversity has also been shown to play a role in determining disease severity and human to human transmission, with “modern” lineages showing a faster progression to disease and shorter latency periods compared to strains from the “ancestral” clades (10-13).

Most of what we know about TB biology today is based on work performed during many decades, most of which has relied on the two canonical reference strains H37Rv and Erdman. Both of these laboratory strains, as well as the clinical strain CDC1551 used by some TB laboratories more recently, belong to MTBC Lineage 4 (14).

H37Rv was first isolated from a patient (H37) with pulmonary tuberculosis in 1905 at the Trudeau Sanatorium in Saranac Lake, New York, while Erdman was isolated from human sputum by William H. Feldman in 1945, at Mayo Clinic, Rochester.

Since its original isolation, H37Rv has been used extensively in biomedical research. The sequence of its genome was published by Cole and colleagues in 1998, which was a breakthrough in TB research (15). Indeed, H37Rv and its genome sequence still provide the backbone for most of TB research today, informing studies ranging from basic biochemistry and microbiology to global omics profiling, systems biology, drug discovery and immunology. However, H37Rv has been passaged countless times in various laboratories, and despite retaining its virulence in mice, it has adapted to laboratory conditions (16). The same is likely true for Erdman and CDC1551 which have been isolated later than H37Rv, but which by now, have also been passaged in the laboratory for several decades. Hence, despite the great progress in our understanding of TB generated through studies based on laboratory strains, there are good reasons to expect that the findings from these studies do not paint the full picture.

Despite the increasing numbers of experimental studies revealing important phenotypic differences across MTBC clinical strains, many of these studies have been difficult to reproduce between different laboratories, and the data are often contradictory. Moreover, linking experimental phenotypes to clinical and epidemiological characteristics of MTBC lineages or strains has been particularly challenging. We propose that part of these challenges could be overcome by standardizing the complement of clinical MTBC strains we study. As a first step, we suggest to broaden the scope of basic and translational TB research by incorporating a set of genetically well-characterized clinical strains representative of the known phylogenetic diversity of the pathogen. In time, the community would accumulate a significant body of data that could support new findings that are more relevant to global TB. To this end it is important that there is a collective agreement to avoid passaging these strains extensively and minimize laboratory adaptation.

Over the years, our group has invested considerable effort into collecting strains from around the world, characterizing them by whole genome sequencing. Our main aim was always to draw evolutionary and phylogeographic inferences (17), however, we also realized the importance of studying this diversity more broadly which is why we used this large strain collection and the associated phylogenomic data to rationally select a subset to be used as reference strains for future research. We believe this set of strains will be of value for the TB research community.

The “MTBC clinical strain reference set” comprises 20 clinical strains covering all 7 known human-adapted MTBC lineages. These strains have been submitted to the Belgian coordinated collections of microorganism (BCCM) strain bank and will be available for anyone interested in the phenotypic impact of MTBC diversity.

## Methods

### Strain selection

We based our initial selection of strains to be included in this reference set on several phylogenetic trees that were built with a combination of genomes from our collection and other public available genomes representing the known global diversity of the human-adapted MTBC. Initially, we picked 43 strains that were intended to represent a diverse sampling of each lineage, comprising several sub-lineages where appropriate. We strove to include strains that were at a basal phylogenetic position within their respective lineage, thus attempting to capture most of the within-lineage diversity. The strains had to be free of known drug resistance mutations and carry only genomic deletions that were congruous with their genetic background, without any rare genomic abnormalities. Moreover, we included strains that were already used in experimental work in the past whenever possible (4, 18).

### Bacterial culture and DNA extraction

All MTBC isolates included into the “MTBC clinical strain reference set”, were processed and were derived from single colonies. Strains were grown in 7H9-Tween 0.05% medium (BD) +/- 40mM sodium pyruvate. We extracted genomic DNA for WGS from cultures in the late exponential phase of growth using the CTAB method (19).

### Spoligotyping

Spoligotyping was performed according to internationally standardized protocols (20). We used KvarQ to derive *in silico* spoligotypes from FASTQ files containing the WGS information (21) when necessary.

### Whole-genome Sequencing

Sequencing libraries were prepared using NEXTERA XT DNA Preparation Kit (Illumina, San Diego, USA). Multiplexed libraries were paired-end sequenced on Illumina HiSeq2500 (Illumina, San Diego, USA) with 151 or 101 cycles at the Genomics Facility Basel.

### Bioinformatic analysis

#### Sequence read alignment and variant determination

The obtained FASTQ files were processed with Trimmomatic v 0.33 (SLIDINGWINDOW: 5:20) (22) to clip Illumina adaptors and trim low quality reads. Any reads shorter than 20 bp were excluded for the downstream analysis. Overlapping paired-end reads were then merged with SeqPrep v 1.2 (overlap size = 15) (https://github.com/jstjohn/SeqPrep). We used BWA v 0.7.13 (mem algorithm) (23) to align the resultant reads to the reconstructed ancestral sequence of MTBC obtained in (24). Duplicated reads were marked by the Mark Duplicates module of Picard v 2.9.1 (https://github.com/broadinstitute/picard) and excluded. The Realigner Target Creator and Indel Realigner modules of GATK v 3.4.0 (25) were used to perform local realignment of reads around indels. To avoid false positive calls Pysam v 0.9.0 (https://github.com/pysam-developers/pysam) was used to exclude reads with alignment score lower than (0.93*read_length)-(read_length*4*0.07)), corresponding to more than 7 miss-matches per 100 bp. SNPs were called with Samtools v 1.2 mpileup (26) and VarScan v 2.4.1 (27) using the following thresholds: minimum mapping quality of 20, minimum base quality at a position of 20, minimum read depth at a position of 7-fold and without strand bias. Only SNPs considered to have reached fixation within a patient were considered (at a within-host frequency of ≥90%). Conversely, when the SNP within-host frequency was ≤10% the ancestor state was called. Additionally, we excluded genomes with average coverage < 15-fold (after all the referred filtering steps). All SNPs were annotated using snpEff v4.11, in accordance with the *M. tuberculosis* H37Rv reference annotation (NC000962). SNPs falling in regions such as PPE and PE-PGRS, phages, insertion sequences and in regions with at least 50 bp identities to other regions in the genome were excluded from the analysis as in (28). Drug resistance-conferring mutations were annotated based on a previously published list (21). Determination of sub-lineage was done using the phylogenetic SNPs according to Stucki *et al*. (28) and to Coll *et al*. (29) (Table 1).

**Table 1.**
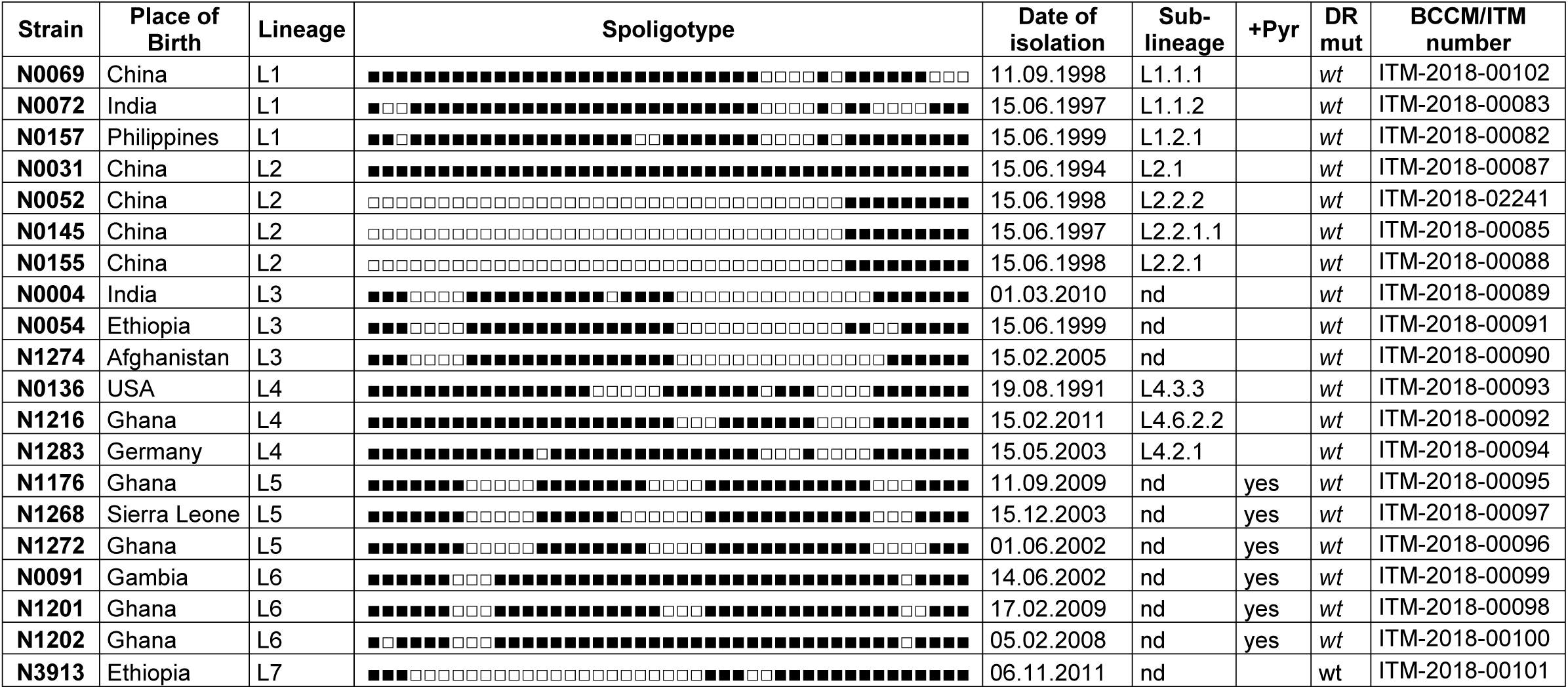
“MTBC clinical strains reference set” list. Place of birth, genotyping data (SNP typing and splogotyping^a^), date of isolation, sublineage classification based on Coll et al. (29), suggested growing conditions and BCCM/ITM number for the strain bank. ^a^Spoligotyping data for each strain are shown, where black squares indicate the presence of a particular spacer and a white square the absence of a particular spacer

#### Phylogenetic reconstruction

To put the reference strain set into a phylogenetic context, we combined them into a phylogeny containing publicly available genomes (n=232) (24). We used all 16.614 variable positions to infer a Maximum Likelihood phylogeny using the MPI parallel version of RAxML (30). We used the GTR model as implemented in RAxML, to perform 1,000 rapid bootstrap inferences, followed by a thorough maximum-likelihood search (30). We show the best-scoring Maximum Likelihood topology. The phylogeny was rooted using *Mycobacterium canettii* as out-group.

## Results and Discussion

### Selection of the “MTBC clinical strain reference set”

Starting from the 43 candidates, we excluded strains that we could not re-grow in the laboratory and those that had any mutation known to confer drug resistance (31). We then selected 20 pan-susceptible clinical strains, based on full genome data for inclusion as reference strains (Table 1). The selected set covers all 7 known phylogenetic lineages of the human-adapted MTBC, but does not include any animal-adapted members of the MTBC. For most lineages, we selected 3 representative strains to maximize the within- and between-lineage diversity. Given the current limited availability of Lineage 7 strains, we were able to include only a single representative. In the case of Lineage 2, we chose 4 strains to cover the wide range of genomic deletions found in this lineage (32). These include N0155, a clinical strain that was used by our group in the past (18), and all the Lineage 2 strains that were transcriptionally profiled by Rose et al. (4). We also included a “Proto-Beijing” strain (N0031), which belongs to a Lineage 2 clade that is phylogenetically basal compared to all other Lineage 2/Beijing strains. Proto-Beijing strains are also characterized by a deletion of RD105 but no deletion in RD207, which differentiates “proto-Beijing” from the regular “Beijing strains (33). Moreover, N0031 shows an ancestral spoligotype with all DR spaces present (Table 1). Its basal branching provides an important contrast to the classical Beijing strains which carry both deletions (33, 34). The Lineage 1 strains N0157 and N0072 have also been transcriptionally profiled (4). In the case of Lineage 4, we selected representative strains of the “generalist” and “specialist” groups as defined by Stucki *et al* (28), respectively N0136 and N1216. The Lineage 4 strain N1283 is a representative of the sub-lineage L4.2 (28), which is also referred to as “Ural” based on spoligotyping (35). The phylogenetic relationships of the 20 reference strains with respect to other representative MTBC strains are shown in Figure 1.

**Figure 1.**
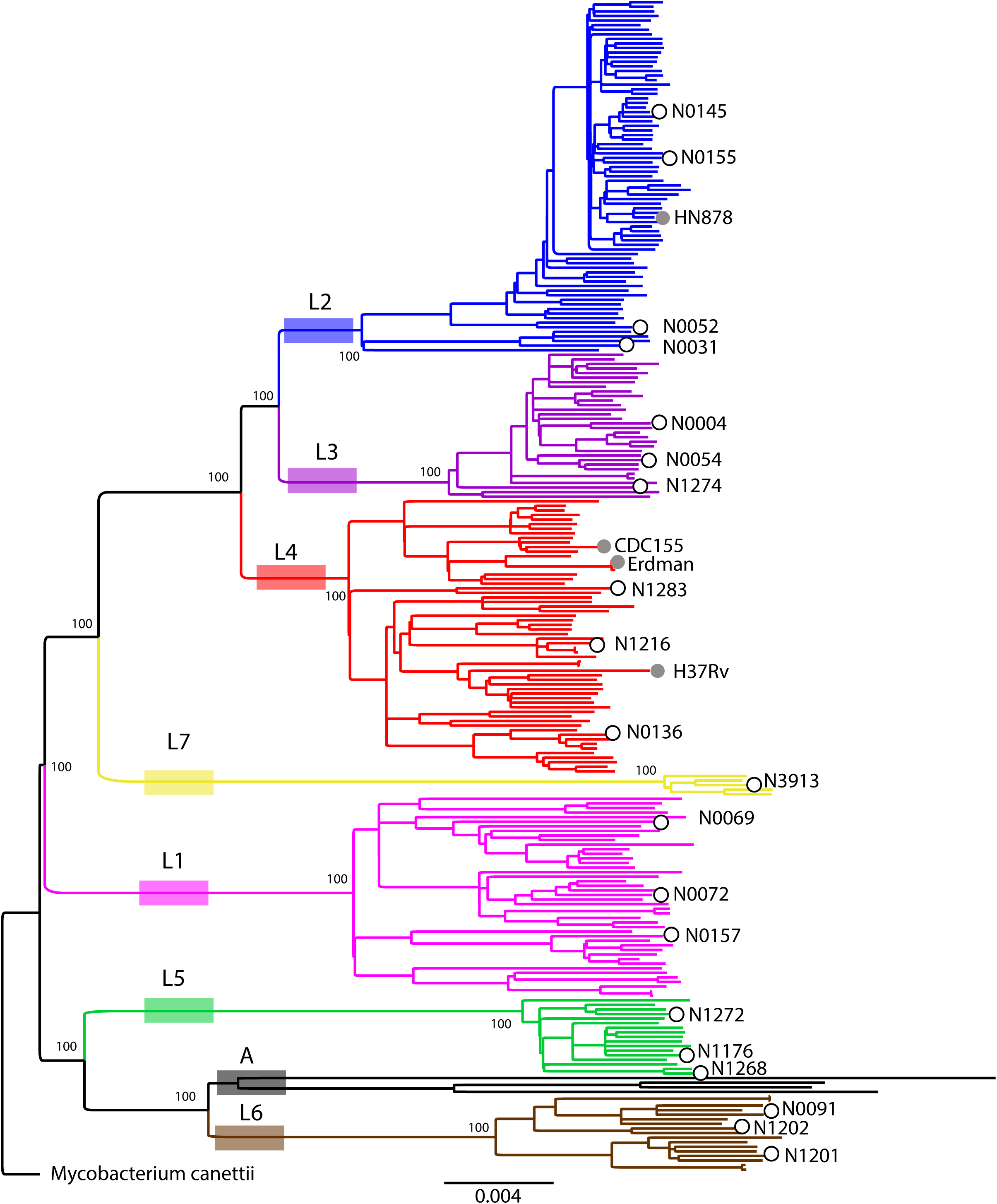
Maximum Likelihood topology of the 20 reference strains (open circles) plus 236 genomes representative of MTBC global diversity. Branch lengths are proportional to nucleotide substitutions and the topology is rooted with *Mycobacterium canettii*. Bootstrap values for clades corresponding to main MTBC lineages are shown. Grey circles indicate the phylogenetic placement of laboratory *M. tuberculosis* strains commonly used.

### Genomic characteristics

All annotated SNPs and insertions/deletions (indels) identified by comparison with the reconstructed MTBC ancestor sequence (36) and considered fixed (at frequency of ≥90%) for each strain are provided as supplementary files. The general characteristics of the genome of each strain are presented in Table 2. We were able to observe all the large genomic deletions reported before (32) as gaps in sequencing coverage. For example, all Lineage 2 strains carried the deletion in RD105 and all but the Proto-Beijing strain also had a deletion in RD207. N0145 and N0155 shared the deletion in RD181, while N0145 harboured an additional deletion in RD150.

**Table 2.** Characteristics of the “MTBC clinical strains reference set” genomes. ^a^ Average read depth after mapping and filtering out duplicated reads. ^b^ Number of SNPs and short Indels considered fixed. ^c^ Percentage of the reference chromosome (H37Rv) to which reads have been mapped. ^d^ Accession Number.

| **Strain** | **Coverage<sup>a</sup>** | **SNPs<sup>b</sup>** | **Indels<sup>b</sup>** | **% Genome Covered<sup>c</sup>** | **AC_Number<sup>d</sup>** |
| --- | --- | --- | --- | --- | --- |
| **N0069** | 81.09 | 898 | 135 | 98.11 | ERR2704679 |
| **N0072** | 72.11 | 894 | 129 | 98.31 | ERR2704680 |
| **N0157** | 74.32 | 894 | 132 | 98.87 | ERR2704704<br>ERR2704685 |
| **N0031** | 66.8 | 845 | 104 | 98.57 | ERR2704676 |
| **N0052** | 110.37 | 862 | 93 | 98.98 | ERR2704677<br>ERR2704699<br>ERR2704698 |
| **N0145** | 39.46 | 875 | 95 | 98.84 | ERR2704702<br>ERR2704701<br>ERR2704683 |
| **N0155** | 115.19 | 897 | 105 | 99.14 | ERR2704703<br>ERR2704684 |
| **N0004** | 46.34 | 873 | 102 | 98.98 | ERR2704698<br>ERR2704675 |
| **N0054** | 64.21 | 886 | 110 | 98.40 | ERR2704678 |
| **N1274** | 80.82 | 874 | 111 | 98.25 | ERR2704693 |
| **N0136** | 52.92 | 823 | 52 | 99.15 | ERR2704682<br>ERR2704700 |
| **N1216** | 66.75 | 817 | 52 | 98.93 | ERR2704705<br>ERR2704689 |
| **N1283** | 52.97 | 831 | 61 | 98.97 | ERR2704709<br>ERR2704694 |
| **N1176** | 76.08 | 934 | 146 | 98.37 | ERR2704686 |
| **N1268** | 51.11 | 937 | 134 | 98.58 | ERR2704706<br>ERR2704690 |
| **N1272** | 73.57 | 908 | 141 | 98.39 | ERR2704708<br>ERR2704707<br>ERR2704692<br>ERR2704691 |
| **N0091** | 72.87 | 1049 | 147 | 98.36 | ERR2704681 |
| **N1201** | 77.64 | 1055 | 148 | 98.39 | ERR2704687 |
| **N1202** | 78.02 | 1015 | 144 | 98.25 | ERR2704688 |
| **N3913** | 100.62 | 1021 | 149 | 99.02 | ERR2704711<br>ERR2704695<br>ERR2704710 |

Given several reports in the literature regarding the importance of the duplication of a part of the genome that includes DosR and DosS (Rv3133 and Rv3134c) for MTBC virulence (4), we included two Lineage 2 strains that carry these duplications – N0031 and N0145. We detected no other large genomic duplications.

### Recommendations for growing and preserving the reference strains

Strains have been deposited in the Belgian Coordinated Collections of Microorganism (BCCM) and can be obtained from the BCCM/ITM http://bccm.belspo.be/services/distribution. Upon receipt, we suggest to grow a large culture of each strain in 7H9 (BD) and freeze multiple glycerol stocks for future use to avoid the acquisition of genetic changes due to laboratory adaptation during sequential sub-culturing (16). Note that some of the strains require that their growth medium to be supplemented with 40mM sodium pyruvate in order improve growth (Table 1).

## Concluding Remarks

For decades, TB research has almost exclusively focused on the two laboratory-adapted MTBC reference strains H37Rv and Erdman. Both strains have provided a common language across TB laboratories allowing knowledge to be built incrementally, with interoperable protocols, results and resources. However, insufficient attention has been given to the fact that both of these strains show patterns of laboratory adaptation and that they do not adequately represent the phylogenetical breadth of the human-adapted MTBC.

The new “MTBC clinical reference set” presented here covers much of this diversity and will provide the TB research community the opportunity to go beyond one single strain/lineage. The potential of sharing and integrating the experimental data generated with this strain set will enrich our understanding of the relationship between genotype and phenotype and potentially lead to fundamental new insights into TB biology. The true impact of genetic diversity in MTBC is slowly coming into focus; however there are still considerable gaps in our understanding. For example, it is known that clinical isolates show variations in drug susceptibility, but the basis for this is unclear. Similarly, the association between drug resistance and specific strain backgrounds has been proposed in several studies, but the underlying mechanism remains unknown. Vaccine and diagnostics development are two areas where understanding the impact of genetic diversity could be key to delivering effective products. Similarly, we are only beginning to scratch the surface of the interplay between bacterial and human genetics at the immune interface. These aspects of MTBC physiology deserve further attention especially due to their potential to have real clinical relevance. At a minimum, testing new TB diagnostics, drugs and vaccines against this strain set will help ensure these innovations are broadly effective.

## Acknowledgements

Computation was performed at sciCORE (http://scicore.unibas.ch/) scientific computing core facility at University of Basel. This work was supported by the Swiss National Science Foundation (grants 310030_166687, IZRJZ3_164171, IZLSZ3_170834 and CRSII5_177163), the European Research Council (309540-EVODRTB) and SystemsX.ch.

